# Antimicrobial Loaded Graft-Copolymer Nanoparticles for Treatment of *Pseudomonas aeruginosa* Infections

**DOI:** 10.1101/2025.07.03.663093

**Authors:** Yadiel Varela Soler, William Xu, Mariana R.N. Lima, Jessica McDonald, Sugeet K. Jagpal, Thomas J. Kirn, Sabiha Hussain, David I. Devore, Charles M. Roth

## Abstract

Nearly 80% of cystic fibrosis patients are affected by persistent lung infections, with *Pseudomonas aeruginosa* being one of the major culprits. Treatment of *P. aeruginosa* is further complicated by its ability to form biofilms. Anionic compounds within the biofilm and thick cystic fibrosis mucus interact with cationic antimicrobials, hindering treatment efficacy. In this study, we investigated the treatment of lung infections by delivering antimicrobials via polyelectrolyte surfactants that are composed of an anionic poly(alkylacrylic acid) backbone with grafted polyetheramine pendent chains. When combined with cationic antimicrobials, they self-assemble into nanoparticles via electrostatic interactions. We assessed the role of backbone chemistry and graft density on nanoparticle physical properties and evaluated the antimicrobial activity of these formulations against planktonic and biofilm cultures of *P. aeruginosa* strains derived from clinical isolates. All synthesized polyelectrolyte surfactants demonstrated high levels of antimicrobial encapsulation, with the extent of drug bound corresponding to the calculated hydrophilic-lipophilic balance values. We observed significantly increased antimicrobial activity against planktonic cultures using nanoformulations containing one of the polyelectrolyte surfactants, PMAA-g-10%J. In contrast, all tested nanoformulations retained, but did not increase, activity against biofilms. By monitoring membrane potentials and nanoparticle uptake, it was found that the nanoparticles directly associate with the bacterial cell membranes, which may enhance drug delivery and underlie the improved activity against the planktonic bacteria. In conclusion, we provide a proof of concept for the design of polyelectrolyte surfactants for the nanoencapsulation and delivery of cationic drug cargoes against *P. aeruginosa* infections.

---

In the United States alone, over 35,000 individuals suffer from cystic fibrosis (CF).^1^ CF is an autosomal, recessive disorder characterized by mutations in the CF transmembrane regulator (CFTR) gene. Defects in the corresponding protein lead to a disruption in the salt-water balance at the tissue surface, ultimately resulting in dehydrated mucus and reduced mucociliary clearance.^2^ The thick mucus allows for proliferation of opportunistic pathogens, such as gram-positive *Staphylococcus aureus* and gram-negative *Pseudomonas aeruginosa* (PA), leading to acute infection or chronic colonization.^3^ Nebulized delivery of cationic aminoglycoside, tobramycin (TB), or the cationic polymyxin, colistin, is the standard of care to treat chronic colonization with PA lung infections in CF patients. Chronic colonization with PA is a leading cause or morbidity and mortality in CF patients.^4^ The use of polymyxin B (PB) for the treatment of PA infections has also increased due to the rise in antimicrobial resistance.^5^ However, the high viscosity and thickness of CF sputum serves as an initial biophysical barrier against antimicrobial treatment, making eradication of organisms such as *Pseudomonas* difficult.^6^

Efficacious treatment of infections using antimicrobials is uniquely challenging due to the formation of bacterial biofilms, which can exhibit 10-1000x lower sensitivity to drug treatments when compared to their planktonic form.^7^ Biofilms are defined as clusters of bacteria that are enveloped in a self-secreted extracellular polymeric substance (EPS) composed of exopolysaccharides, extracellular DNA, lipids, and proteins.^8^ Negatively charged components of the EPS are known to interact with cationic drugs, effectively hindering their diffusion across the biofilm and acting as an additional barrier to antimicrobial treatment.^9^

Nanoparticles (NPs) provide a valuable opportunity to overcome current therapeutic limitations. They can confer a number of advantageous properties, including the potential to control drug release^10-13^ and to protect the drug from degradation.^11, 14-16^ The success of the FDA approved liposomal amikacin nanoformulation, Arikayce®, for the treatment of lung infections has further motivated the study of new drug delivery systems in this area.^17^ A variety of polymer chemistries are available to impart desirable drug delivery properties to their formed nanoparticles. For instance, the incorporation of polyethylene glycol (PEG) chains to the NP surface has been shown to enhance penetration in cystic fibrosis mucus.^13, 18-25^ By encapsulating the cationic drugs within NPs, electrostatic interactions with anionic compounds of the mucus can potentially be reduced, enhancing diffusion. For this purpose, we have developed polyelectrolyte surfactants capable of encapsulating cationic drugs via electrostatic self-assembly.^26-29^ These surface-active polymers are composed of poly(alkylacrylic acid) backbones grafted with polyetheramine pendent chains (Jeffamine® M-2070) via carbodiimide coupling, as shown by the reaction schematic in **Figure 1**. Amphiphilic chains, such as those offered by Jeffamine® M-2070, are capable of improving particle stability, protecting against the reticuloendothelial system,^26^ and enhancing membrane penetration.^13^

**Figure 1.**
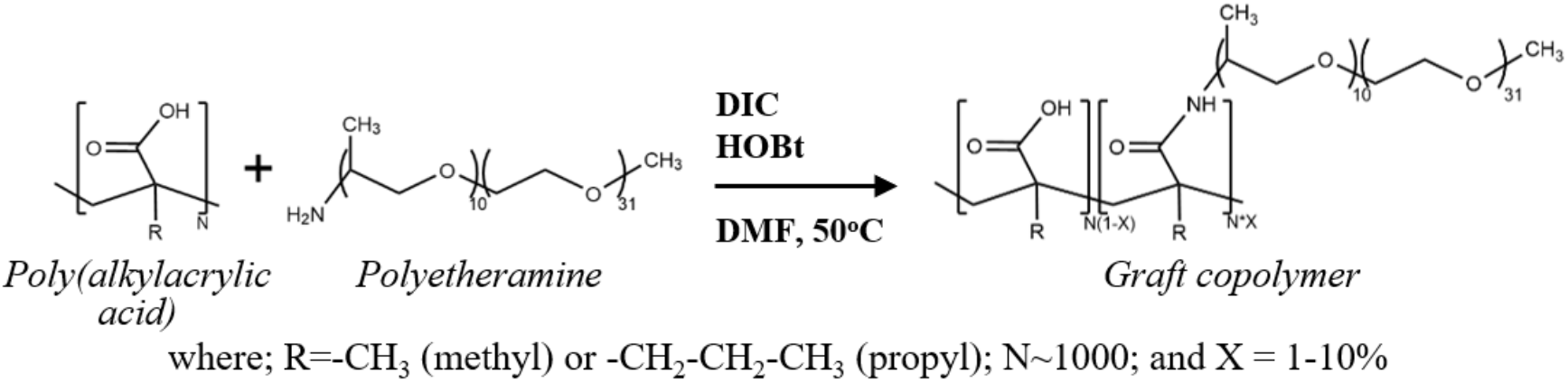
Schematic representation of graft copolymer synthesis. Polyetheramine chains (Jeffamine® M-2070) were grafted onto poly(alkylacrylic acid) backbones (poly(methylacrylic acid), PMAA and poly(propylacrylic acid), PPAA) via carbodiimide coupling reaction.^27^

In this work, we developed a small library of graft copolymer NPs based on the design shown in **Figure 1**. We demonstrated that they can be loaded with cationic antimicrobials, TB and PB, and we evaluated the influence of polymer chemistry on drug loading and release as well as on nanoparticle size. The fine-tunable features of these graft copolymers, backbone chemistry and graft density, allowed for control of NP size, release and loading capacity.^27^ Furthermore, we assessed the activity of the resulting nanocomplexes against CF strains of PA in both planktonic and biofilm states.

## RESULTS

### Graft Copolymer Synthesis and Nanoparticle Characterization

Starting with the design of a backbone poly(alkylacrylic acid) grafted with a polyetheramine, as shown in **Figure 1**, we performed calculations of the hydrophilic-lipophilic balance (HLB) using group contribution methods for various possible configurations (**Table 1**).^30-33^ We selected poly(methacrylic acid) (PMAA) and poly(propylacrylic acid) (PPAA) as the backbone polyelectrolytes based on their previously demonstrated ability to form nanoparticles with cationic peptides and lipids,^28, 34^ while allowing us to probe different levels of backbone hydrophobicity. The levels of graft density evaluated were chosen based on graft copolymer solubility and previous studies.^34^ Due to the lipophilicity of the propyl group in PPAA, polymer constructs with PMAA backbones possess higher HLB values than those with PPAA. While the addition of polyetheramine chains slightly increases polymer hydrophilicity, HLB values are primarily driven by backbone chemistry.

**Table 1.** HLBs of select polyelectrolyte surfactants.

| Polymer | HLB (calculated) |
| --- | --- |
| PMAA | 7.68 |
| PMAA-g-1%J | 7.79 |
| PMAA-g-5%J | 8.27 |
| PMAA-g-10%J | 8.85 |
| PPAA | 6.72 |
| PPAA-g-1%J | 6.85 |
| PPAA-g-5%J | 7.36 |
| PPAA-g-10%J | 8.00 |

*Note:* HLB calculations based on the fully protonated form.

Both PMAA and PPAA were successfully grafted with nominally 10% Jeffamine® M-2070 (J) by carbodiimide coupling. While the graft copolymer PPAA-g-10%J was successfully synthesized, it was not soluble in aqueous solution and therefore not pursued further. The actual graft densities were determined by NMR analysis using the ratio of Jeffamine -CH_3_ to methacrylic acid -CH_2_-groups (**Figure S1**). In all cases, the experimentally determined graft densities were at least 80% of the targeted values (**Table 2**).

**Table 2.** Measured graft density (GD) of PS and the hydrodynamic diameter of unloaded and loaded PS nanoparticles.

| PS | GD [%] | Diameter [nm] |  |  |
| --- | --- | --- | --- | --- |
|  |  | No Drug | Tobramycin | Polymyxin B |
| PMAA-g-10%J | 8.3 | 71 ± 7 | 167 ± 6 | 219 ± 6 |
| PMAA-g-5%J | 4.0 | 89 ± 4 | 160 ± 3 | 197 ± 4 |
| PPAA-g-5%J | 5.3 | 160 ± 6 | 234 ± 37 | 204 ± 3 |
| PPAA-g-1%J | 1.0 | 98 ± 2 | 173 ± 13 | 160 ± 3 |

*Note.* Nanoparticles were prepared at a charge ratio (+/-) of 0.5. Values of hydrodynamic diameter represent the mean ± std. dev.

All four graft copolymers successfully encapsulated the cationic antimicrobials, tobramycin (TB) and polymyxin B (PB), due to favorable electrostatic interactions between the cationic groups of the drugs and the anionic backbone of the graft copolymer.^29, 35^ In the absence of drug, each graft copolymer self-assembled to form nanoparticles of approximately 100 nm in size. When combined with cationic antimicrobials, prepared at a positive to negative charge ratio of 0.5 based on previous studies,^34, 36^ the particle size increased by an amount in the range of 50-200% depending on the particular combination of drug and graft copolymer (**Table 2**). For polymers with a PMAA backbone, PB-loaded NPs showed larger sizes than those with TB. Graft density had a significant impact on NP size for PPAA-based formulations, with sizes being greater at higher graft densities. This trend was also observed in PMAA-based formulations, but to a lesser extent. At the same target graft percentage of 5%, PPAA based formulations demonstrated significantly greater NP sizes than PMAA based formulations. Other than PPAA-g-5%J+PB formulations, NPs were found to be stable, as no significant change in their size distribution was detected after 4 weeks of storage at 4°C (**Figure S2**).

Nanoparticles were evaluated for their binding efficiency and release kinetics. All formulations, other than TB loaded PPAA-g-1%, bound drug at efficiencies above 90% (**Figure 2**). PMAA-based formulations exhibited statistically stronger binding than those with a PPAA backbone. This trend correlates with the HLB values for the PSs (**Table 1**), which is indicative of the importance of polar group interactions between the PS and the drugs in the drug binding process. The TB release rate varied markedly with graft copolymer composition, with faster release being observed for the more weakly bound formulations (**Figure 2B**). PB exhibited slow release from all NP formulations, making it difficult to discern a trend with regard to graft copolymer composition (**Figure 2C**).

**Figure 2.**
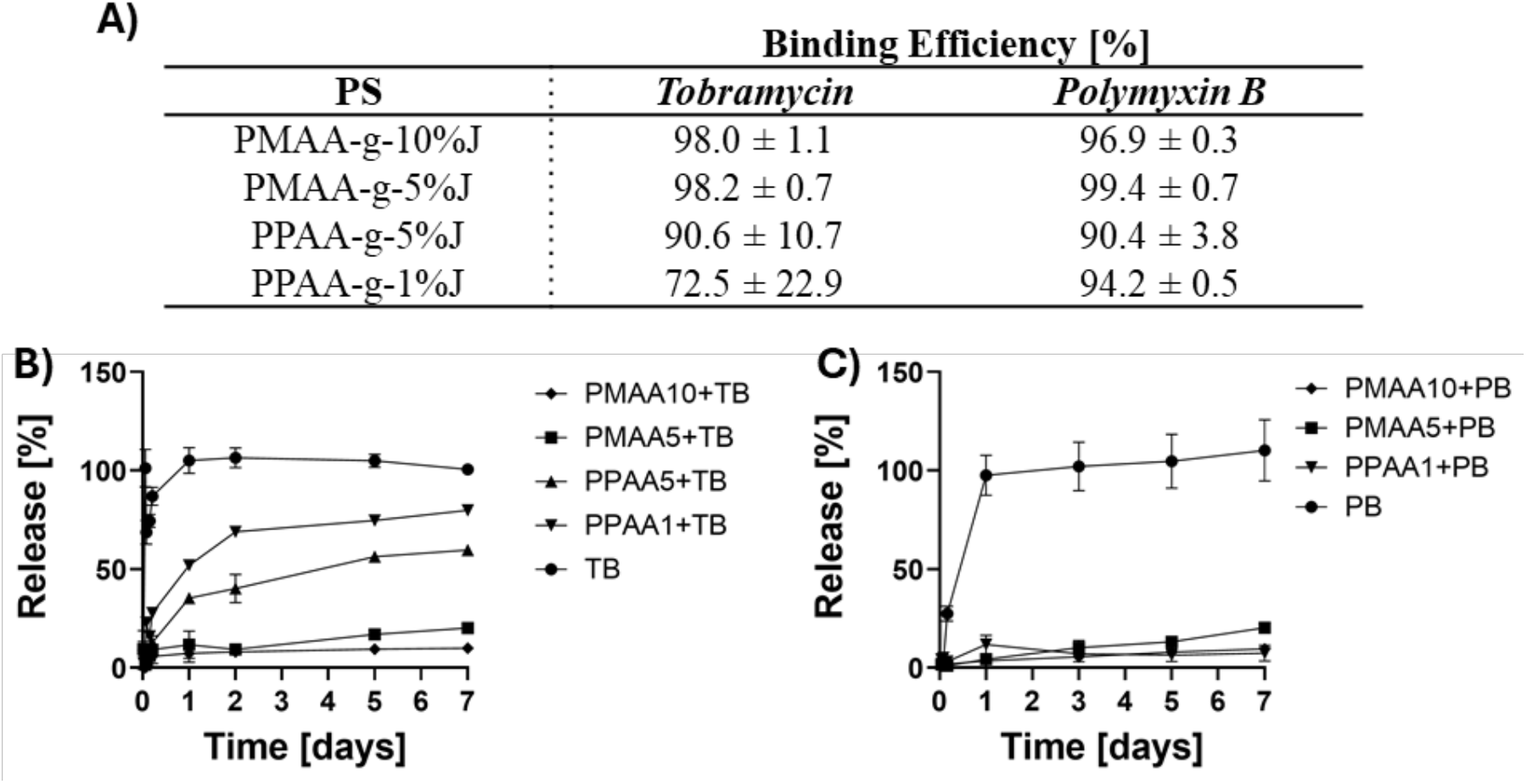
Drug-polymer interaction at an initial charge ratio of 0.5. **A)** Initial binding efficiency; values represent mean ± std. dev. Release kinetics of free and encapsulated **B)** TB and **C)** PB in phosphate buffered saline. Initial concentrations of NPs were 150 μg/mL and 300 μg/mL for TB and PB, respectively.

### Growth Inhibition of Planktonic *P. aeruginosa*

Having established favorable conditions for encapsulation of cationic antimicrobials into nanoparticles, we sought to evaluate how nanoformulation affects the drugs’ antimicrobial properties. To determine the minimum inhibitory concentration (MIC) required for growth inhibition of planktonic cultures, four clinical strains of PA from CF patients were cultured for 24 hours with varying concentrations of TB-NPs, TB, PB-NPs, and PB, according to microbroth dilution assay protocols.^37^ Modest variations in MICs were observed across the set of clinical strains, reflecting that each strain came from an individual patient source with unique clinical and drug treatment history (**Figure 3**). PMAA based formulations demonstrated a statistically significant, minimum 2-fold reduction of the MICs relative to the free drugs. Meanwhile, formulations made with PPAA retained the same level of activity as the drug alone. The most effective formulations for both TB and PB were those made with PMAA-g-10%J.

**Figure 3.**
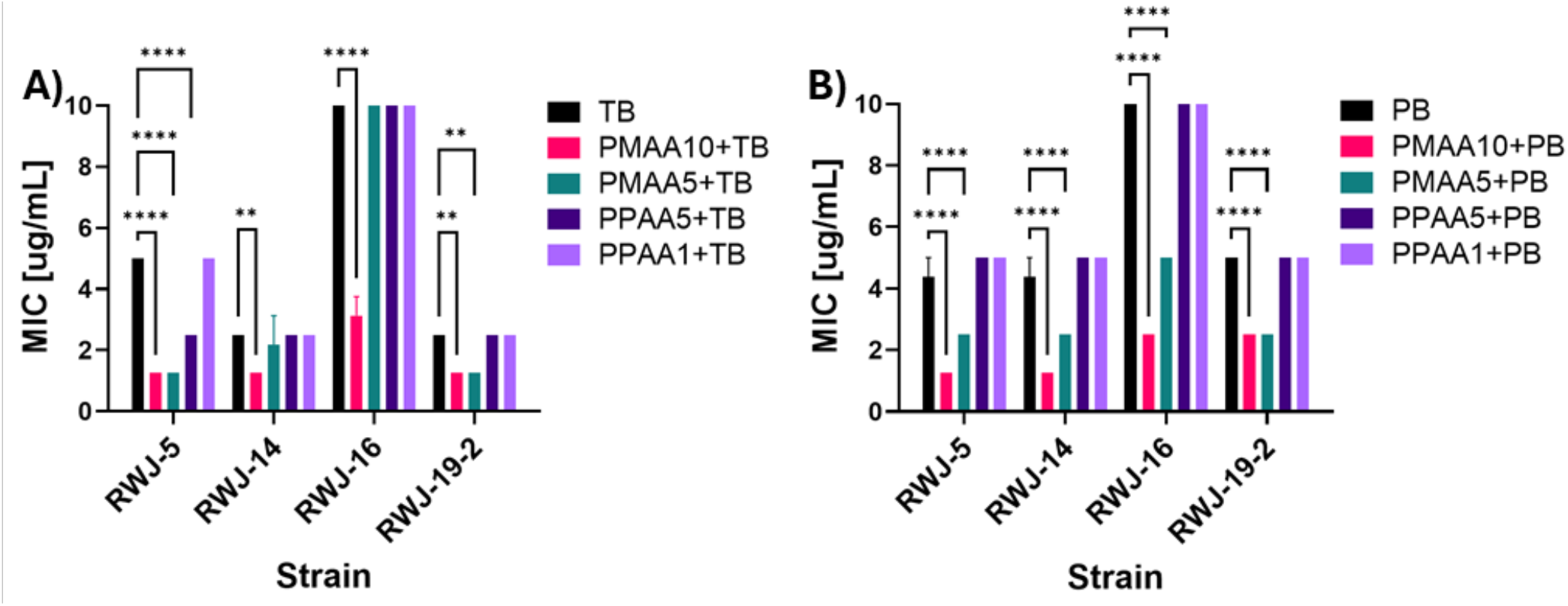
Minimum inhibitory concentration of **A)** TB and **B)** PB nanoparticles against *Pseudomonas aeruginosa* clinical strains. Clinical strains of PA were incubated with decreasing concentrations of NP formulations for 24 hours. The MIC was determined to be the lowest concentration to inhibit 95% of PA growth.

### Eradication of PA Biofilms

Biofilm related lung infections affect over 75% of CF patients,^38^ making biofilms an important bacterial state against which to test the antimicrobial activity of our NP formulations. The minimum biofilm eradication concentration (MBEC) was determined as the lowest concentration required to eradicate 95% of an already established biofilm.^39^ Compared to growth inhibition (MIC) assays, MBEC values across seven clinical strains of PA exhibited greater variation; therefore, this data is displayed on a logarithmic axis (**Figure 4**). The MBEC assays show that, on average, NP formulations eradicate already established biofilms to roughly the same extent as free drug (**Figure 4**). Strain- and drug-specific differences were also observed in the impact of nanoformulation, with decreased MBEC values (increased effectiveness) observed in TB-based nanoformulations against 2 of 7 PA strains. Nonetheless, no overall trends were observed in the MBECs of the different strains after being exposed to the various formulations. The graft-copolymer treatment alone showed no reduction in biofilm growth (data not shown).

**Figure 4.**
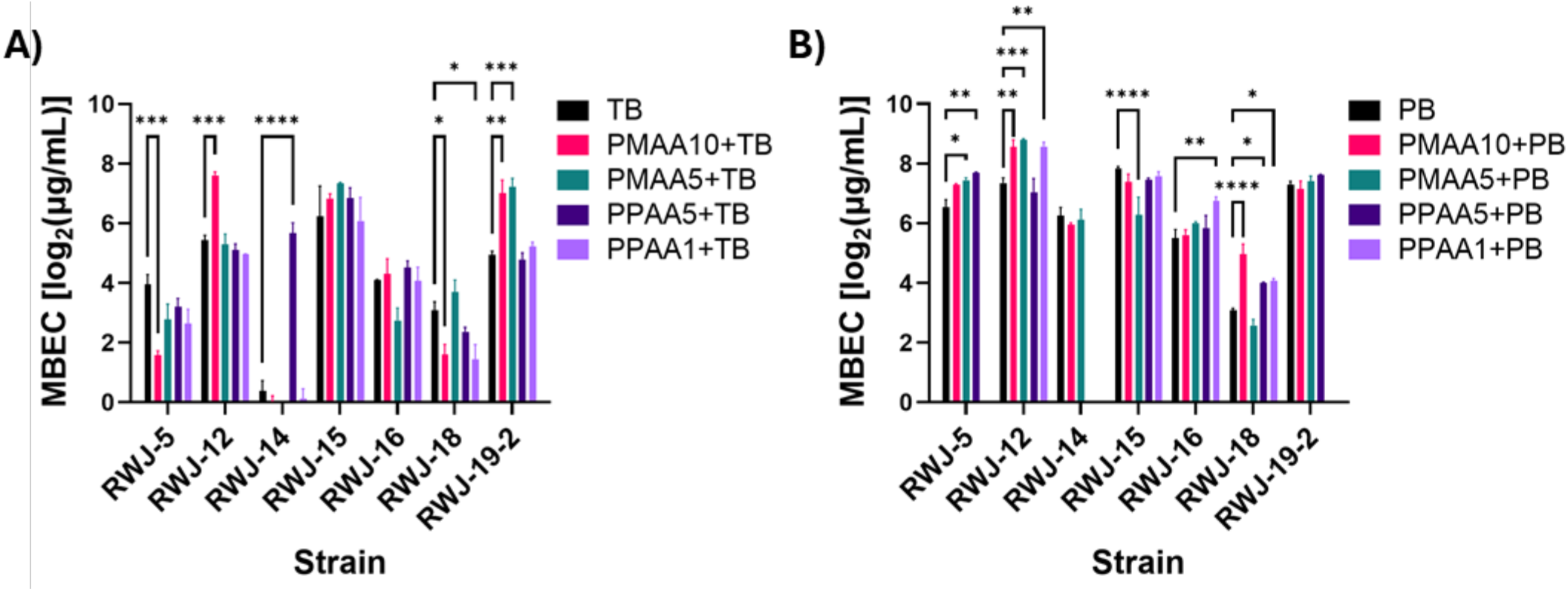
Minimum biofilm eradication concentration of **A)** TB and **B)** PB nanoparticles against *Pseudomonas aeruginosa* clinical strains. Biofilms were grown on the surfaces of 96-well plates by incubating clinical strains of PA for 48 hours. They were then exposed to a dilution series of concentrations of NP formulations for 24 hours. The MBEC was determined via curve fitting as the concentration where 95% of the biofilm was eradicated and is displayed on a log_2_ scale due to the large variation in values across the strains.

### Exploring NP Interactions with Bacteria

Based on the observed increase in antimicrobial activity of PMAA-based formulations against planktonic cells, we sought to assess NP interaction with the bacterial cells more directly. Using Cy5.5 fluorescently tagged PMAA-g-10%J, denoted as Pol-Cy, the extent of interaction with bacteria was evaluated. NPs prepared with Cy5.5-labeled polymer increased approximately 15% in size compared to the unlabeled NPs (**Figure S3**). Bacteria in the mid-log phase were exposed to NPs at their MIC and incubated for 1 hour. After incubation, bacteria were separated from the media by centrifugation, the levels of free Pol-Cy were measured in the supernatant, and the amount associated with bacteria was calculated by mass balance. Approximately 20% of both TB and PB loaded NPs were found to be associated with bacteria within the timeframe of the experiment (**Figure 5A**). Polymer alone showed no sign of uptake by cells.

**Figure 5.**
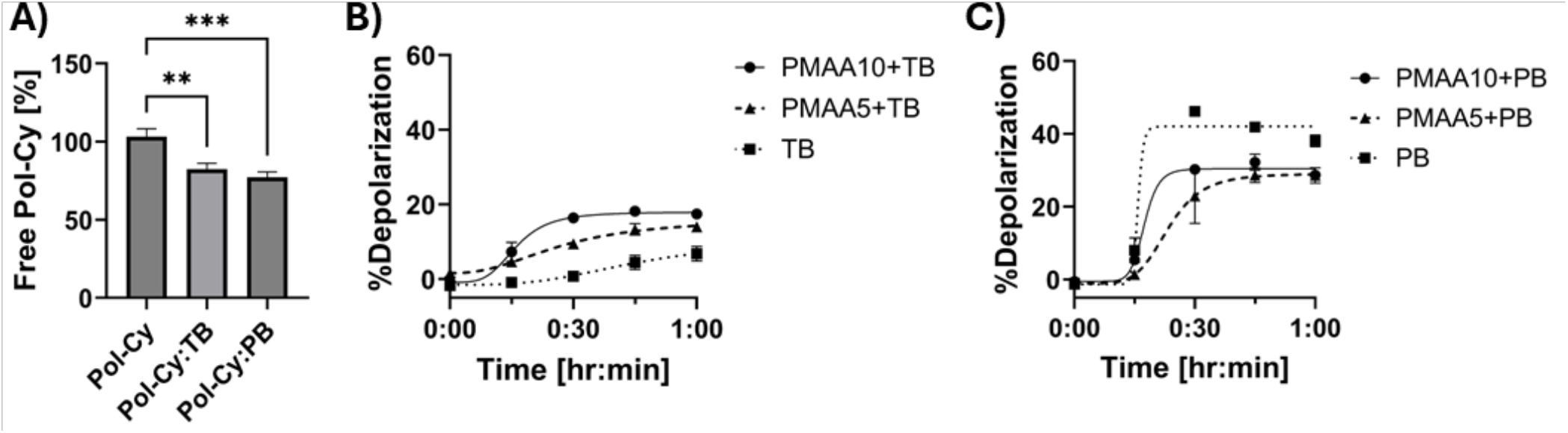
Evaluating the interaction between NPs and bacteria. **A)** Relative amount of free Pol-Cy in media following a one-hour incubation at the free drug MIC. The relative depolarization of bacteria membrane following addition of **B)** TB and **C)** PB formulations.

To further explore the interaction between NPs and bacteria, the planktonic cells were incubated with the membrane potential-sensitive fluorescent probe DiSC_3_(5). This probe localizes in the membrane of bacteria and is released upon membrane depolarization, leading to an increase in fluorescence. Upon exposing the planktonic bacteria to the NP formulations, membrane depolarization was measured over time as seen in **Figure 5B-C**. PB alone produced significantly greater membrane depolarization than TB alone. This was consistent with the known antimicrobial mechanism of TB, which does not involve membrane disruption, and PB, which strongly disrupts the membranes.^40, 41^ When encapsulated in nanoformulations, the PS-TB nanoparticles exhibited increased membrane depolarization (**Figure 5B**), while the PS-PB nanoparticles produced less membrane depolarization than PB alone (**Figure 5C**). In effect, the PS enhanced depolarization caused by TB and mitigated the depolarization caused by PB.

## DISCUSSION

The pathophysiological state of the lungs in cystic fibrosis patients poses a serious limitation to current therapies for treating the *P. aeruginosa* infections that frequently take hold in the CF population. The thick CF mucus and overproduction of extracellular DNA hinder diffusion of cationic antimicrobials. Furthermore, PA possesses multiple mechanisms to resist the entry and effect of antimicrobial drugs, including through the formation of biofilms, which pose an additional biophysical barrier to drug transport. We designed partially anionic graft copolymers capable of encapsulating cationic antimicrobials to form nanoparticles via electrostatic interactions. Formulation into nanoparticles enables controlled release and alters the interaction of cationic cargoes with bacterial cell membranes. Depending on their surface characteristics, nanoparticles have the potential to penetrate the physiological microenvironments of mucus and biofilm as well.

The polyelectrolyte surfactants utilized in this work were designed to produce anionic, amphiphilic nanoparticles. The balance between hydrophilic and lipophilic groups is determined by the composition of both the backbone and the graft. The carboxylic acid of the alkylacrylic acid backbone provides a titratable anionic charge, while the identity of the alkyl moiety provides a means to tune lipophilicity, with methyl and propyl variants studied here. The polyetheramine graft utilized in this study comprises a random copolymer of ethylene oxide (EO) and propylene oxide (PO) groups, under the trade name Jeffamine®. The triblock copolymers of EO and PO, under the trade name Pluronics®, have been formulated and used by others for delivery of a variety of drugs, including antibiotics to treat bacterial infections in both planktonic and biofilm states.^42-45^ The polyetheramine grafts in our copolymer can be tuned by their composition (EO/PO ratio), molecular weight, and graft density. In this work, we focused on Jeffamine® M-2070, which has 31 EO groups and 10 PO groups, resulting in a molecular weight of approximately 2,000 Da. We have previously developed grafts utilizing Jeffamine® M-2005, with 6 EO and 20 PO. This highly lipophilic variant was insoluble in aqueous media when the graft density exceeded 1% and proved hemolytic,^28^ motivating the current composition with its moderate proportion of PO groups. As a means to quantify the various polyelectrolyte surfactants, which vary in backbone type and Jeffamine® graft density, we employed HLB calculations, based on group contribution methods.^30, 31^ The HLB calculations reveal that backbone chemistry is the primary contributor to polyelectrolyte surfactant lipophilicity, with graft density playing a secondary role (**Table 1**). The propyl group in the PPAA backbone leads to a significant increase in hydrophobicity relative to PMAA. The addition of Jeffamine® M-2070 chains results in a small increase in polymer hydrophilicity (HLB), due to the greater number of hydrophilic ethylene oxide groups compared to the number of lipophilic propylene oxide groups.

The sizes, binding, and release of these nanocomplexes are governed by an interplay of electrostatic interactions, steric hindrance, hydrogen bonding, and hydrophobic interactions between the polymer backbone and the drug. In the absence of any drug, the amphiphilic nature of the polyelectrolytes causes them to self-assemble into nanoparticles that are generally under 100 nm in diameter (**Table 2**). In all cases, NP diameters increased when either tobramycin or polymyxin B was formulated into the nanoparticles. For a given backbone, NPs with a higher graft density showed increased sizes, which could be partly attributable to a decrease in the strength of electrostatic interactions. As the graft density increases, fewer backbone carboxylic groups become available, leading to less densely packed NPs of greater diameter. Furthermore, polymers with a high graft density are bulkier, producing steric hindrance and increasing particle size. In addition to electrostatic interactions between the backbone carboxyl and drug amine groups, both also possess numerous groups capable of hydrogen bonding with each other and surrounding water molecules. These forces are in competition with hydrophobic interactions generated by the polymer backbone, especially for the more hydrophobic PPAA backbone. While the PPAA backbone’s increased hydrophobicity could promote tighter association of the NP core, the relatively larger size of the propyl group, relative to a methyl group, could explain the larger size observed in PPAA nanoparticles. Alternatively, the more greatly hydrophobic core promotes the formation of nanoparticles with a greater number of polymer molecules, thus increasing particle size. The increase in size with increasing graft density was less drastic for PB loaded NPs, which could suggest that the hydrophobic tail of PB slightly stabilizes these interactions.

Polymer variations in binding and release were present to a greater extent in nanoparticles formed with the small molecule aminoglycoside, tobramycin (TB), than in those formed with the lipopeptide, polymyxin B (PB). The PMAA-based polyelectrolytes exhibited both greater binding and slower release of TB than their PPAA-based counterparts. In addition, higher graft density polyelectrolytes tended to have greater binding and slower release. The trends with respect to polyelectrolyte backbone and graft are consistent with the trends in calculated HLB values, as is the greater influence of backbone over graft density. For polymyxin B, on the other hand, high extents of binding and slow release were observed for all the polyelectrolytes tested. The difference in binding and release characteristics between the two drugs can be attributed to their structure. The more polar drug, TB, is primarily trapped by electrostatic interactions; meanwhile, the fatty acid tail of PB confers a second mechanism for interaction with the backbone and amphiphilic chains of the graft copolymers.

Gram-negative bacteria are characterized by their outer membrane, which comprises phospholipids, lipopolysaccharides, and pendent oligosaccharides. These collectively present a range of lipophilic and hydrophilic barriers that might be traversed best by an amphiphilic structure. We utilized fluorescence localization and membrane depolarization assays to probe the interactions between amphiphilic nanoparticles formed from the PMAA-backbone polymers, which were the most biologically active, and the well-characterized PAO1 strain of Gram-negative *Pseudomonas aeruginosa*. Unloaded nanoparticles showed no uptake by bacteria, suggesting that the negative net charge of the polyelectrolyte in the absence of cationic cargo prevented the nanoparticles from overcoming repulsion by the net negative surface charge of the bacterial membrane. When loaded with cationic drugs, TB and PB, cellular localization was observed, indicating membrane interaction or uptake had taken place. This interaction can be attributed to the presence of “free” cationic charges on the bound drugs; that is, not every cationic amine group of the two poly-cationic drugs was condensed on (i.e., electrostatically coupled with) a backbone carboxylic acid group and those “free” cationic groups could then electrostatically bind with anionic groups on the bacterial cells’ membrane surfaces.

By probing membrane potential, we were able to further evaluate these interactions. PB is known to act by directly disrupting the outer membrane, leading to cell death. Consistent with this mechanism, the greatest depolarization was observed with PB alone, followed closely by NP formulations. PB is known to behave like a biosurfactant, binding to phospholipids in the bacterial membrane and leading to its disruption.^46-48^ On the other hand, TB acts by blocking mRNA translation and alone did not lead to significant disruption of the membrane potential. When encapsulated, significantly greater membrane depolarization was observed, suggesting that our polyelectrolyte surfactant nanoparticles promote membrane interaction. Graft density seemed to play a role in membrane interaction, with NPs created from a more highly grafted polymer, PMAA-g-10%J, showing slightly increased depolarization. Considering its nearly equal HLB value to PMAA-g-5%J, it is likely that lipophilicity does not play a role. Rather, the significantly greater amount of polyetheramine chains likely causes a more drastic impact on membrane fluidity and permeability.^49-51^ Combined, these results suggest that polyelectrolyte surfactant nanoparticles promote interaction with bacterial membranes and cellular entry, which could explain the increased antimicrobial activity observed in MICs.

While PMAA-based formulations showed a 2-fold or greater increase in antimicrobial activity against planktonic PA, their effectiveness against already established PA biofilms was shown to be highly strain- and formulation-specific. Previous researchers have observed strain differences regarding the ability of surfactants to inhibit biofilm formation and suggested that this behavior is related to the expression of bacterial lipases.^52^ A similar mechanism could explain the significant strain variations observed in this work. It is likely that further optimization of formulation conditions could provide more robust activity against PA biofilms. For example, the nanoparticles utilized in most of these studies were formulated at a charge ratio (+/-) of 0.5. We did observe improved activity against biofilms when the charge ratio of PS-TB nanoparticles was increased to 1.0 (**Figure S4**). However, those nanoparticles were electrostatically neutral and proved to be colloidally unstable; as a result, they were not pursued further.^36^ It is likely that further optimization towards the dual objectives of biofilm activity and colloidal stability can be performed by varying formulation conditions and/or polyelectrolyte surfactant chemistry.

In this work, we explored variation of backbone chemistry and graft density in polyelectrolyte surfactants as design features for the encapsulation and delivery of clinically essential, commercial cationic antimicrobials active against gram-negative bacteria. The results were consistent with our prior investigation of PS nanoparticle complexes of an experimental, biodegradable cationic polypeptide antimicrobial against gram-positive bacteria.^26^ Nanoparticles made with PMAA-based graft copolymers provided the greatest benefit against planktonic bacteria and modest benefit against biofilms from certain PA strains. Formulations containing the graft copolymer PMAA-g-10%J, which carries the highest HLB value among the tested polyelectrolyte surfactants, exhibited high drug binding, sustained release, membrane depolarization, and increased microbiological activity against clinical strains of *P. aeruginosa*. Using HLB as a metric to guide design, next generation polyelectrolyte surfactants with different graft modifications may offer greater activity against fully formed biofilms. Combined with their capacity to control release and their optimal delivery size, PMAA-based graft copolymers offer a promising delivery system for increasing effectiveness against *Pseudomonas aeruginosa* infections.

## MATERIALS AND METHODS

### Reagents and General Materials

Reagents were purchased from commercial sources unless otherwise noted. Poly(methylacrylic acid) (PMAA) and poly(propylacrylic acid) (PPAA) were purchased from Polysciences, Inc. (Warrington, PA) and Polymer Source (Dorval, Quebec, Canada), respectively. Hydroxybenzotriazole (HOBt), N,N-diisopropylcarbodiimide (DIPC), dimethylformamide (DMF), 1-Ethyl-3-(3′-dimethylaminopropyl)carbodiimide (EDC), N-Hydroxysuccinimide (NHS), 3,3-dipropylthiadicarbocyanine iodide (DiSC_3_(5)), sodium hydroxide (NaOH) and dimethyl sulfoxide (DMSO) were purchased from MilliporeSigma (Burlington, MA), along with the cationic drugs tobramycin (TB) and polymyxin B (PB) sulfate. Jeffamine® M-2070 was purchased from Huntsman Corp (Woodlands, TX). 2,4-dinitrofluorobenzene was purchased from Chem-Impex (Wood Dale, IL). Acetonitrile (ACN) and Tris(hydroxymethyl) aminomethane (Tris Base) were purchased from Fisher Scientific (Waltham, MA). Cy5.5-amine was purchased from Broadpharm (San Diego, CA). Bacterial cultures were grown in Trypticase™ Soy Broth from BD (Franklin Lakes, NJ). Bioavailability was determined with Alamar Blue staining, purchased from Bio-Rad (Philadelphia, PA).

Slide-A-Lyzer dialysis cassettes (3 mL, 10kDa MWCO) were purchased from Thermo Scientific (Waltham, MA). Syringe filters were obtained from VWR (Radnor, PA). Sensoplate glass bottom 96-well plates used for size measurements were purchased from Greiner Bio-One (Monroe, NC). 30 kDa MWCO Microcon® centrifuge filters were purchased from MilliporeSigma (Burlington, MA). For microtiter plate assays with incubations over 24 hr, plates were sealed with Rayon Films from VWR® (Radnor, PA).

### Bacterial strains

The acquisition of bacterial strains was approved by the Rutgers University Institutional Review Board under protocol #Pro20170001066 as non-human subjects research, as no identifying information was associated with the samples. During spring 2019, sputum samples were collected routinely from patients at the Rutgers-Robert Wood Johnson Cystic Fibrosis Center as part of their clinical care. When samples tested positive for *Pseudomonas aeruginosa*, the strain was isolated and subcultured on blood agar plates. After storage, each strain was grown on tryptic soy agar (TSA) plates. Those that remained viable after storage and reinoculation were used for antimicrobial studies.

### Instrumentation

Graft density was determined by proton nuclear magnetic resonance (^1^H-NMR) analysis on a 500 MHz-Varian Inova (Santa Clara, CA) spectrometer. Size distribution of NPs was determined by dynamic light scattering (DLS) on a Wyatt DynaPro® Plate Reader III (Santa Barbara, CA). Absorbance and fluorescence measurements were performed on an Infinite M200 Pro Microplate Reader from Tecan (Männedorf, Switzerland). Drug concentrations were evaluated using an Agilent 1260 Infinity II High-Performance Liquid Chromatography (HPLC) System (Santa Clara, CA).

### Hydrophilic-lipophilic balance (HLB) calculations

The surface activity of surfactants, including those comprised of amphiphilic graft copolymers, can be characterized by their hydrophile:lipophile balance (HLB). HLB values were calculated using a semi-empirical method based on **equation (1)** developed by Davies:^30-32^

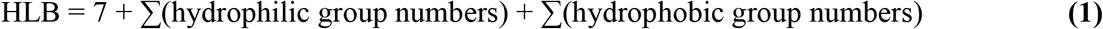

The pertinent group numbers^30-33^ for the PSs are summarized in **Table 3**:

**Table 3.** HLB Group Numbers

|  |  |
| --- | --- |
| -(EO)<sub>n</sub>- | 0.33n, where n= degree of polymerization |
| -(PO)<sub>n</sub>- | - 0.15n, where n= degree of polymerization |
| -CH<sub>x</sub>- | - 0.475, where x=1,2,3 |
| -COOH | 2.1 |
| -COO<sup>-</sup> Na<sup>+</sup> | 19.1 |
| -CONH- | 3.7 |

### Graft Copolymer Synthesis

For PPAA based graft copolymers, PPAA (0.44 mol of repeat units) was combined with HOBt (0.044 mmol) and DIPC (0.044 mmol) and dissolved in 5 mL of THF. After 45 min of mixing, a 10% excess of Jeffamine® based on the targeted graft densities was added and mixed further for 30 hours at 50°C. Graft copolymer was dried by roto-evaporation and resuspended in a mixture of 10%v/v 1 N NaOH and 90%v/v PBS. Polymer suspension was then dialyzed in water for 3 days using a 10 kDa MWCO Slide-A-Lyzer, with three bath changes in the first 6 hours and two every day after. The dialyzed solution was collected from within the cassettes, lyophilized and characterized by ^1^H-NMR using deuterated DMSO as a solvent. The degree of labeling was determined as the ratio of the (-CH_2_-) peak of PPAA and (-CH_3_) peak of Jeffamine® M-2070.

Similarly, for PMAA-based graft copolymers, PMAA (2.32 mmol of repeat units), HOBt (0.232 mmol), and a 3% excess of the corresponding amount of Jeffamine® M-2070 (0.116, 0.232 mmol for 5, 10 mol%, respectively) were dissolved in 2.5 mL of DMF. DIPC (0.232 mmol) was then added to the mixture and stirred for 30 hours at 50°C. DMF was removed by roto-evaporation and the graft copolymer resuspended in methanol. The graft copolymer was then precipitated by drop-wise transfer to 20 mL of diethyl ether. A Buchner filter with fritted disk was used to filter the product which was then allowed to air dry for 1 hour. The graft copolymer was then placed under vacuum for a final 24 hours. Proton NMR was used to characterize actual grafting density by determining the ratio of PMAA (-CH_2_) and Jeffamine® M-2070 (-CH_3_).

A small amount of PMAA-g-10%J was also grafted with Cy5.5, in order to allow fluorescent tracking. Briefly, PMAA-g-10%J (1 mmol) was combined with EDC (50 mmol), NHS (200 mmol), Cy5.5-amine (0.06 mmol), and 10mL of DMSO. These were allowed to react for 24 hours at room temperature, covered from light. Polymer purification and characterization were carried out as previously explained.

### Nanoparticle Preparation

First, 24 mg of lyophilized graft-copolymers were dissolved overnight in 4 mL of 0.1 M NaOH (diluted from 10M NaOH, Sigma-Aldrich, St. Louis, MO). Graft-copolymers were then transferred to 3 mL 10kDa MWCO Slide-A-Lyzer from Thermo Scientific (Waltham, MA) and dialyzed in Milli-Q water for 4 days with bath changes at 0.5-, 1-, 3-, 6 hours, and at 1-, 2-, and 3 days. After dialysis was complete, volume was determined, and necessary amounts of polymer were collected for NP formation. The remaining graft copolymer was stored at 4°C under dark.

Meanwhile, stock TB and PB solutions were prepared in Milli-Q water at 2.5 mg/mL and 5 mg/mL, respectively. NP formation was initiated by transferring 4 mL of dialyzed graft copolymer to a 15 mL glass scintillation vial and sonicating for 20 min in an ice bath. A stir bar was placed in the vial, and the contents were mixed vigorously on ice. Based on the tatget grafting density and polymer concentration, the necessary amount of drug to obtain a charge ratio (CR) of 0.5 was calculated and slowly (∼1 drop / 5 seconds) added to the vial containing the graft-copolymer solution. The mixture was stirred rapidly for an additional 10 minutes in the ice bath. The mixture was sonicated in an ice bath for an additional 20 minutes, after which the NPs were diluted to the concentration desired for experiments.

### Nanoparticle Characterization: Size, Stability, Binding, Release and Degradation

The size of the NPs was determined by dynamic light scattering (DLS) using the Wyatt DynaPro® Plate Reader III (Santa Barbara, CA). A 500 μL volume of NPs was filtered using 0.45 μm pore size, 13 mm diameter nylon syringe filters. From the filtered NP formulations, 105 μL were transferred to a glass bottom 96-well plate and their size was measured. The stability of the NPs was evaluated by monitoring their particle size over time when stored at 4°C.

The binding efficiency of the various graft-copolymers was determined by placing 500 μL of NPs in 30 kDa MWCO Centrifuge filters and centrifuging at 14,000g for two cycles of twelve minutes. The TB and PB concentrations in the filtrate were determined using an Agilent 1260 Infinity II high performance liquid chromatography (HPLC) system with the operating parameters shown in **Table 4**. TB determination by HPLC required its derivatization with 2,4-dinitrofluorobenzene (DNFB). To derivatize TB, 400 μL of sample were combined in a 5 mL volumetric flask with 1 mL of 10 mg/mL DNFB in methanol and 1 mL of 3 mg/mL tris(hydroxymethyl) aminomethane in 80%v/v dimethyl sulfoxide. The mixture was incubated for 50 min at 60°C in a water bath and was then cooled for 10 min, after which the contents were diluted with acetonitrile.^53^ Samples were filtered with 0.45 μm pore size, 13 mm diameter nylon syringe filters, after which 2 mL were transferred to HPLC vials.

**Table 4.** Chromatographic conditions.^53, 54^

|  | **TB** | **PB** |
| --- | --- | --- |
| **Column:** | Agilent Eclipse XDB-C18,<br>4.6x150 mm, 5 μm, 80 A | Agilent Zorbax Extend-C18<br>250 x 4.6 mm, 5 μm, 80 A |
| **Column Temp:** | 25°C | 30°C |
| **Flow Rate:** | 1.2 mL/min | 1.0 mL/min |
| **Wavelength:** | 365 nm | 215 nm |
| **Injection Vol:** | 20 μL | 100 μL |
| **Mobile Phase** | 59% ACN – 1% 1N H<sub>2</sub>SO<sub>4</sub> –<br>40% 1g/L Tris Base | 20% ACN – 80% 4.5g/L<br>Na<sub>2</sub>SO<sub>4</sub>, pH 2.3 |

The drugs’ release rate from NPs was determined by placing 2 mL of NPs in 10 kDa MWCO Slide-A-Lyzer cassettes. The cassettes were placed in a beaker containing 120 mL of PBS. Samples from the reservoir were collected at select timepoints, and the total volume of PBS was replenished after each sample collection. Drug concentration was determined by HPLC.

### Minimum Inhibitory Concentration (MIC)

Seven PA clinical strains collected from CF patients in the Robert Wood Johnson Hospital and the commercially available PAO1 strain were cultured overnight from frozen glycerol stocks at 37°C and 215 rpm in Trypticase™ Soy Broth (TSB).^37^ PA cultures were maintained by performing 1:20 dilutions into TSB. For microbial assays, the bacterial load was determined by measuring the optical density at 600 nm (OD_600_) and diluting to 1×10^7^ CFU/mL. A volume of 100 μL of the diluted culture was transferred to a 96-well plate, followed by 100 μL of treatment groups and controls to their respective wells. Treatment concentrations for TB and PB (and their corresponding NPs) started at 34 μg/mL and 64 μg/mL, respectively, followed by 2-fold serial dilutions. All treatment groups were performed in triplicate. The plates were incubated at 37°C overnight, after which they were imaged. The MIC was defined as the lowest concentration of drug capable of preventing visible bacterial growth.

### Minimum Biofilm Eradication Concentration (MBEC)

PA biofilms were grown by incubating 100 μL of a 1×10^7^ CFU/mL culture in 96-well plates at 37°C for 48 hours. The plates were sealed with rayon films. After 48 hours, the breathable film was removed and media containing unbound bacteria was carefully pipetted out of each well so as not disturb the biofilms. To each well, 100 μL of the corresponding treatment group and controls were added. Drug concentrations for formulations of TB and PB started at 256 μg/mL and 512 μg/mL respectively, followed by 2-fold serial dilutions. Plates were resealed with a breathable film and incubated for 24 hr at 37°C. After the second incubation, the film was removed, and the wells were emptied. To each well, 100 μL of a 10% Alamar Blue solution (prepared by diluting in TSB) was added and allowed to incubate for 1-2 hours at 37°C. Finally, to quantify cell viability, fluorescence was measured with an Infinite M200 Pro Microplate Reader (Ex. 530 nm, Em. 590 nm). The MBEC was defined as the concentration at which 95% of the biofilm was eradicated and was determined from the raw fluorescence data using curve fitting in a MATLAB code.

### Membrane Interaction and NP Uptake

PAO1 was diluted to an OD_600_ of 0.5 in LB media and spiked with the corresponding formulation for a concentration equivalent to the MIC. NPs used for this experiment were prepared using Cy5.5 labeled PMAA-g-10%J. The bacteria suspensions were incubated at 37°C and 150 rpm for 1 hour and then centrifuged at 10,000 rcf for 5 minutes. A total of 100 μL of supernatant was transferred to a 96-well plate and the fluorescence measured (Ex. 683 nm / Em. 703 nm) using a plate reader.

### Membrane Potential Assay

For membrane potential analysis, an overnight culture of PAO1 was diluted to an OD_600_ of 0.5 and washed three times with 5 mM HEPES, pH 7.2 buffer. Bacteria were then pelleted and resuspended in a buffer composed of 5 mM HEPES, 20 mM glucose, 0.2 mM EDTA, 0.1 M KCl, and 0.4 μM DiSC3(5).^55^ The cell suspension was then incubated for 30 min at 37°C. In a 96-well plate, 100 μL of cell suspension were combined with 100 μL of dye solution. The fluorescence (Ex. 622 nm / Em. 670 nm) was immediately monitored every 15 min for 1 hour using a microplate reader. The positive control was 1% Triton X-100, and every condition was evaluated in triplicate.

### Statistical Analysis

All experiments were performed in triplicate, unless otherwise indicated. Plots and statistical analyses were performed using GraphPad Prism 10.4. All data are shown as mean ± standard error of the mean (SEM) unless otherwise specified. Statistical tests were performed using a significance level of p<0.05. ANOVAs were performed along with multiple comparison tests to identify statistical differences between groups.

## ASSOCIATED CONTENT

### Supporting Information

The Supporting Information is available free of charge. Representative ^1^H NMR spectra for synthesized PMAA-g-10%J can be found in **Figure S1**, plots showing nanoparticles size over time are provided in **Figure S2**, bar graphs representing size of nanoparticles prepared with cy5.5 tagged PMAA-g-10%J in **Figure S3**, and graphs showing the minimum biofilm eradication concentration of formulations prepared at a charge ratio of 1, **Figure S4**.

## AUTHOR INFORMATION

### Author Contributions

Conceptualization: D.D., C.R.; resources: S.J., T.K., S.H.; methodology and investigation: Y.V., M.L., W.X, J.M.; visualization: Y.V., W.X.; supervision: D.D., C.R.; project administration: C.R.; writing–original draft: Y.V.; writing–review and editing: D.D, C.R. All authors have given approval to the final version of the manuscript.

### Notes

The authors declare no competing financial interest.

## ACKNOWLEDGMENT

Previous iterations of this project were undertaken by several lab members including Dr. Ritu Goyal, Jamal Keyes, Pooja Patel, Youssef Mohamed, and Michael Holloway.

## ABBREVIATIONS USED

PS: Polyelectrolyte Surfactant
NP: nanoparticle
PA: Pseudomonas aeruginosa
TB: tobramycin
PB: polymyxin B
CF: cystic fibrosis
PMAA: poly(methacrylic acid)
PPAA: poly(propylacrylic acid)
J: Jeffamine M-2070

