## Supporting Information for "Antimicrobial Loaded Graft-Copolymer Nanoparticles for Treatment of *Pseudomonas aeruginosa* Infections"

### NMR Spectrum of Synthesized Polymer

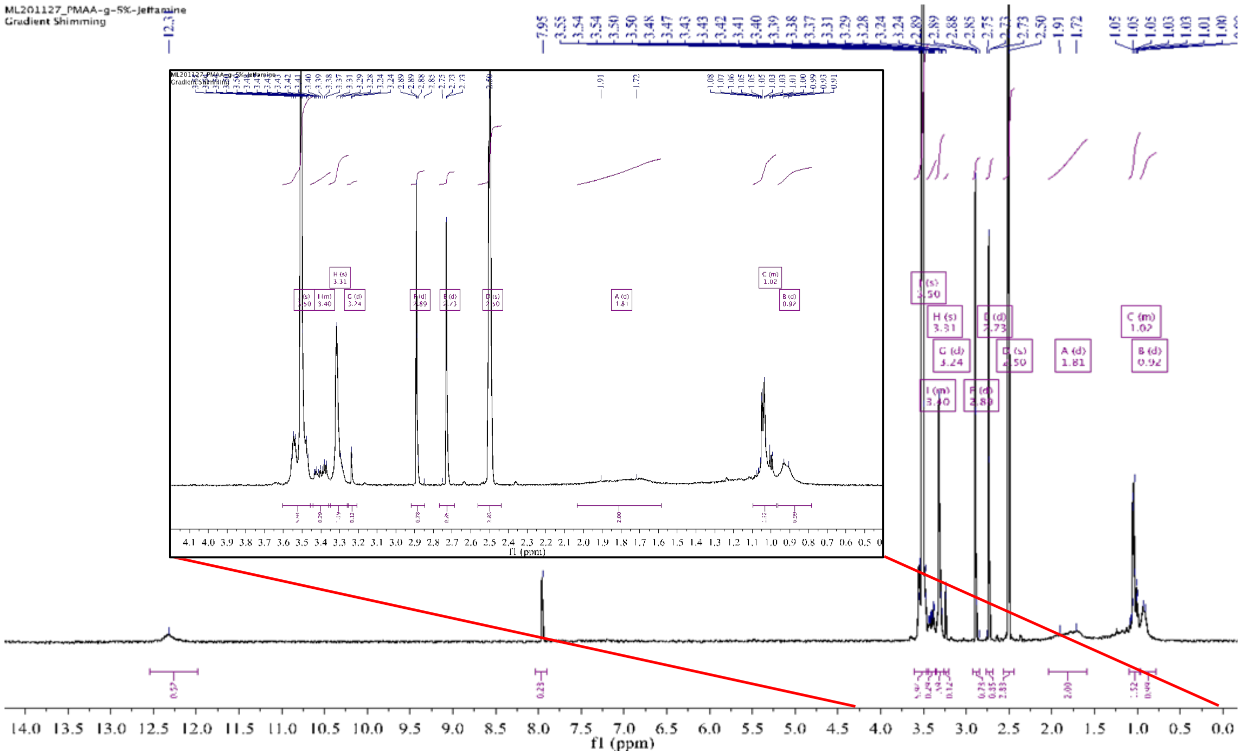


**Figure S1.** Sample H-NMR of PMAA-g-5%J used to determine graft density with chemical shifts (ppm) at 1.81 for PMAA and 3.24 for Jeffamine® M-2070.

**Stability of NPs Over Time at 4$^{o}$C**

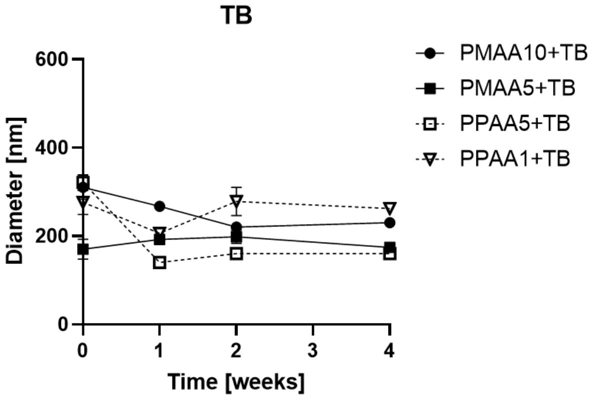


**Figure S2.** Diameter of TB and PB formulations over time.

**Size of NPs when prepared with Cy5.5-labeled PMAA-g-10%J**

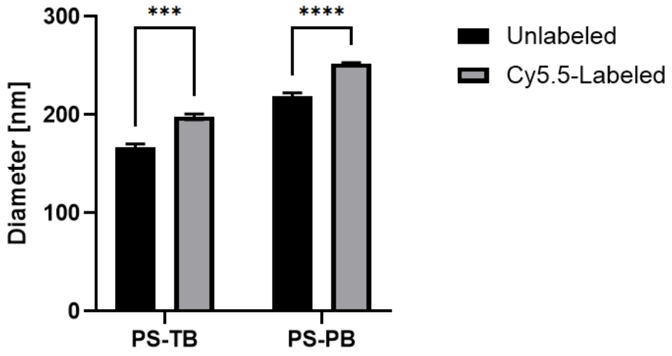


**Figure S3.** Hydrodynamic diameter of TB and PB formulations when prepared with Cy5.5-labeled PMAA-g-10%J.

**Minimum Biofilm Eradication Concentration (MBEC)**

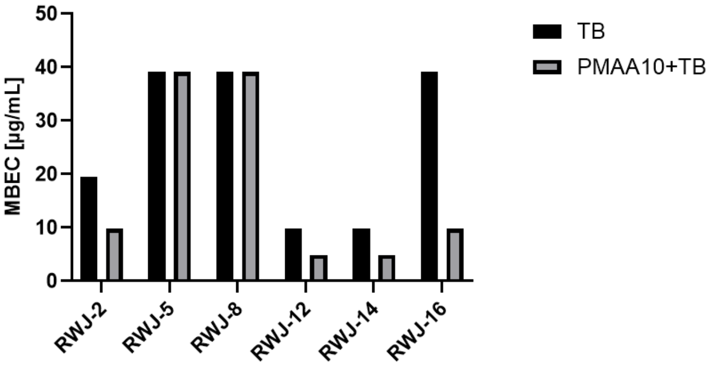


**Figure S4.** MBEC for different strains of *P. aeruginosa* from clinical isolates when using free TB or NPs with PMAA-g-10%J at a charge ratio of 1.[1]
